# Modeling the impact of *ex-nido* transmitted parasites on ant colony dynamics

**DOI:** 10.1101/470575

**Authors:** Lauren E. Quevillon, David P. Hughes, Jessica M. Conway

## Abstract

Infectious disease outbreaks are a common constraint of group living organisms. Ants (Hymenoptera: Formicidae) live in large, dense colonies and are host to a diverse range of parasites and pathogens, facilitating the possibility of epidemic-induced collapse. However, the majority of parasites infecting ants require a period of development outside of the nest before they can transmit to their next ant host (‘*ex-nido*’ transmission) and the impact of these parasites on colony dynamics is unknown. Here we develop a mathematical model to assess ant colony dynamics in the presence of such parasites. We find that under field-realistic model conditions, such parasites are unlikely to cause the epidemic collapse of mature ant colonies, unless colony birth rate drops below 0.2328 ants/day. The preponderance of *ex-nido* transmitting parasites infecting ants and their limited epidemiological impact on colony dynamics may partly explain why collapsed ant colonies are rarely, if ever, observed in natural populations.

## Introduction

Infectious diseases capable of massive mortality events appear to be an unavoidable consequence of living in large, complex societies. As humans evolved from small groups of hunter-gatherers into larger agrarian societies and then cities, we have seen an increase in the number of outbreaks that our societies experience (1, 2). Some diseases (i.e. measles, seasonal influenza) are ‘crowd diseases’ that can only be maintained because of large numbers of individuals living in close proximity. Outbreaks readily occur in large natural and managed animal populations as well. Boom-bust cycles in Gypsy moths driven by virus epidemics (3), Ebola and anthrax outbreaks amongst chimpanzees and gorillas (4, 5), and recurrent foot-and-mouth disease epizootics in commercial bovid all demonstrate that high density, group living organisms often contend with the steep costs of intense disease burden.

However, infectious disease outbreaks may not necessarily be an unavoidable consequence of evolving to live in dense groups. Social insects in the order Hymenoptera (ants, bees, and wasps), and the ants in particular, exemplify the living conditions that exaggerate the perceived risk of infectious disease spread. These animal groups are defined by eusociality, the highest form of social organization, in which there is a reproductive division of labor, overlapping generations, and cooperative brood care (6). Eusociality means that these groups often live in extremely dense living conditions with large colony sizes (up to millions of individuals; (7), reviewed in (8)), and have an average higher genetic relatedness compared to other animal groups. This risk is further compounded by nesting habitats in soil and decaying wood that put colonies at increased risk of contact with microbial loads capable of causing disease (9). Additionally, neighboring colonies overlap in their foraging ecology and compete for the same food resources, enhancing the potential for inter-colony transmission (10, 11). Indeed, social insects are host to a diverse range of pathogens, parasites, and parasitoids (hereafter ‘parasites’) (12, 13). However, ants and other social insects have achieved incredible ecological success (14, 15), and effectively managing their disease burden could be one underlying reason for their continued success over evolutionary time.

Attempting to understand how social insects have contended with considerable parasite pressure over their long evolutionary history has driven empirical research during the past two decades. This work has uncovered many disease-fighting mechanisms that social insects have in their arsenal, including physiological, behavioral, and organizational defenses at both the individual- and colony-levels (reviewed in (12, 16–19)). Some mechanisms, such as allogrooming (the grooming of nest mates) (20, 21) and the transfer of immune modulators or antibiotic secretions between nest mates (21, 22), are evolutionary innovations following their transition to eusociality, but it is likely that many of these immune mechanisms were present prior to the transition to eusociality (23).

Complementary theoretical studies have used a variety of modeling frameworks to test the relative efficacies of these mechanisms against parasites that are capable of direct transmission from one infected nest mate to another. For example, deterministic compartmental modeling has been used to assess differences in epidemiological outcomes when passive versus active immune modulators are passed between nest mates (24), and how grooming, a behavioral defense used to remove infectious particles prior to infection, could impact epidemic potential inside colonies (25). Others have used individual-based modeling approaches to investigate how interaction heterogeneity (26), nest architecture (27), and immunity and hygienic behavior (28) impact disease severity and colony survival. These modeling approaches have been very useful for comparing the relative efficacies of antidisease defenses against parasites that are capable of transmission inside the nest.

One key assumption underlying most of the above work is that parasites infecting ants are capable of direct, ant-to-ant transmission inside the nest. However, the majority of parasite species that infect ants have lifecycles that require a period of development outside of the colony (‘*ex-nido*’), often as free-living stages or in other host species, before they are able to infect new hosts (Fig. 1) (12, 13). The amount of time such parasites need before they are capable of infecting a new ant host varies considerably, from days to months or even years. For example, following emergence from their ant host, phorid flies must mate and then find new hosts to oviposit into within days (29). Trematodes infecting ants must complete development inside their vertebrate final host, mate and release eggs to be consumed by a snail before another ant can be infected, a process which can take months (30). Finally, in temperate systems, the zombie-ant fungus *Ophiocordyceps* does not develop the sexual stages needed to release spores and infect new ants until many months following the death of its original ant host (31). Thus, with these ‘*ex-nido*’ parasites, direct transmission between nest mates inside colonies is not possible, potentially precluding the threat of major disease outbreaks inside colonies.

**Fig. 1.**
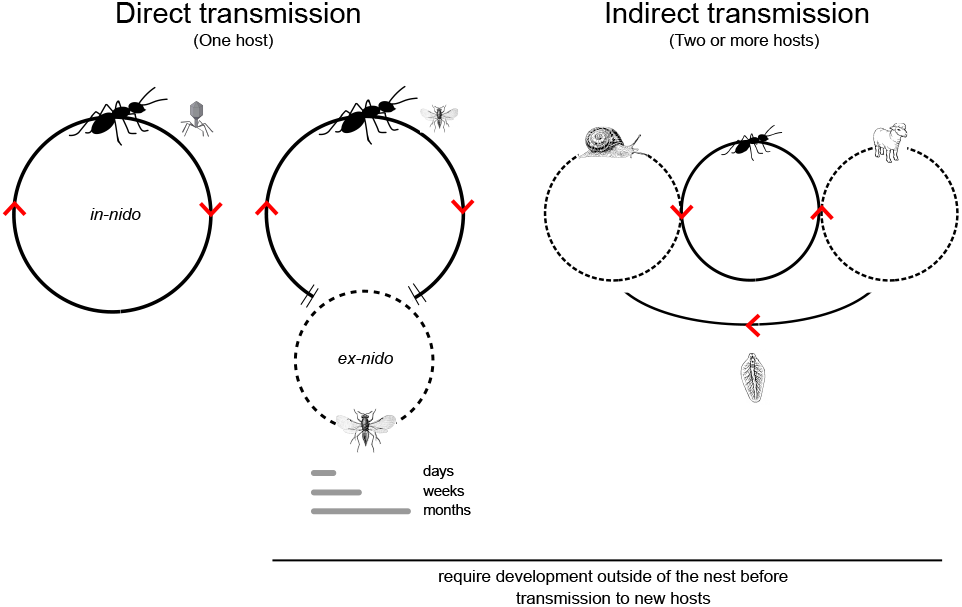
Transmission strategies used by parasites infecting ants. (a) Direct, *in-nido* parasitism occurs when ants are the only host infected and transmission can readily occur from ant-to-ant inside the nest. Examples include bacteria, viruses, and some entomopathogenic fungi. (b) Direct, *ex-nido* parasitism occurs when ants are the only host infected, but the parasite needs to complete development in the extranidal environment before it is capable of transmitting to the next ant host. Examples include parasitoid wasps and flies. (c) Indirect, *ex-nido* transmission occurs when more than one host species is required for the parasite to complete its lifecycle. The use of multiple hosts necessitates development outside of the nest. Examples include cestode, trematode, and some nematode worms.

To date, no theoretical studies have investigated the potential impact of *ex-nido* parasites on ant colony dynamics. Here, we formally explore the consequences of *ex-nido* parasites by building a model of ant colony dynamics in the absence of parasite pressure; we then extend it to a ‘susceptible-infected-removed’ (SIR)-type model for the disease dynamics of parasites transmitted outside of ant colonies. While this model was built to capture ant colony dynamics, it is generally applicable to other social insect colonies. We explore how changing colony birth and parasite transmission rates impact colony dynamics and show that for a range of biological plausible parameter values, colony collapse is unlikely to occur. Finally we explore how other parameters (i.e. proportion of the colony foraging, parasite developmental rate) impact the potential impact of *ex-nido* parasitism on ant colony dynamics. Our work suggests that disease dynamics in ant and other social insect societies, in which *ex-nido* parasitism predominates rather than parasites directly transmitting between nest mates, are fundamentally different from other social living organisms like mammals, which may partly explain the enduring ecological success of the social insects.

## Methods

### A. Model of ant colony dynamics in the presence and absence of *ex-nido* transmitting parasites

Our model of mature ant colony dynamics in the presence and absence of *ex-nido* parasites is given by Eqs. (1-4) and summarized schematically in Fig. 2. In the absence of parasitism, we make the simplifying assumption that there are three developmental and sociological castes (compartments) that individuals transition through in ant colonies, listed below:

- **Brood (B)**: a period of biological development during which individuals transition from egg through several larval stages.
- **Nest workers (N)**: a variable period of time during which individuals participate in intranidal tasks such as brood care and nest maintenance.
- **Foragers (F_*s*_)**: a variable period of time that older ants transition into according to colony need; foraging ants gather food and participate in extranidal territory defense and maintenance, which puts them at greater risk for both predation and parasitism.

We believe that this captures sufficient biological realism of mature ant colony functioning without over-complicating the model. While some ant species do have other specialized castes (e.g. soldiers in army ants, repletes in honey pot ants, etc.), the basic colony demographic structure we have selected applies broadly to the majority of ant colonies (10, 32). Our investigation of colony dynamics in the presence of *ex-nido* parasites, i.e. parasites that require a developmental period outside of the nest before transmission to new hosts, is motivated by the biology of the most prevalent type of ant-infecting parasite that we find from our extensive review of the literature (13), parasite records in (12, 29, 33–37), among many others). We model a parasite that encounters and infects a forager ant in the extranidal (outside of the nest) environment, causes the mortality of that ant after some developmental period, and ultimately transmits to its next host in the extranidal environment (i.e. no direct ant-to-ant transmission within the nest). Thus, our model of colony dynamics in the presence of parasitism includes a fourth compartment, **infected foragers (F**_*i*_). When the force of infection term *β* (described in detail below) is zero, Eqs. (1 - 3) reduce to the uninfected model, which is used to validate parameter choice and the form of the model. When the force of infection term *β* is greater than zero, Eqs. (1 - 4) represent the infected model.

**Fig. 2.**
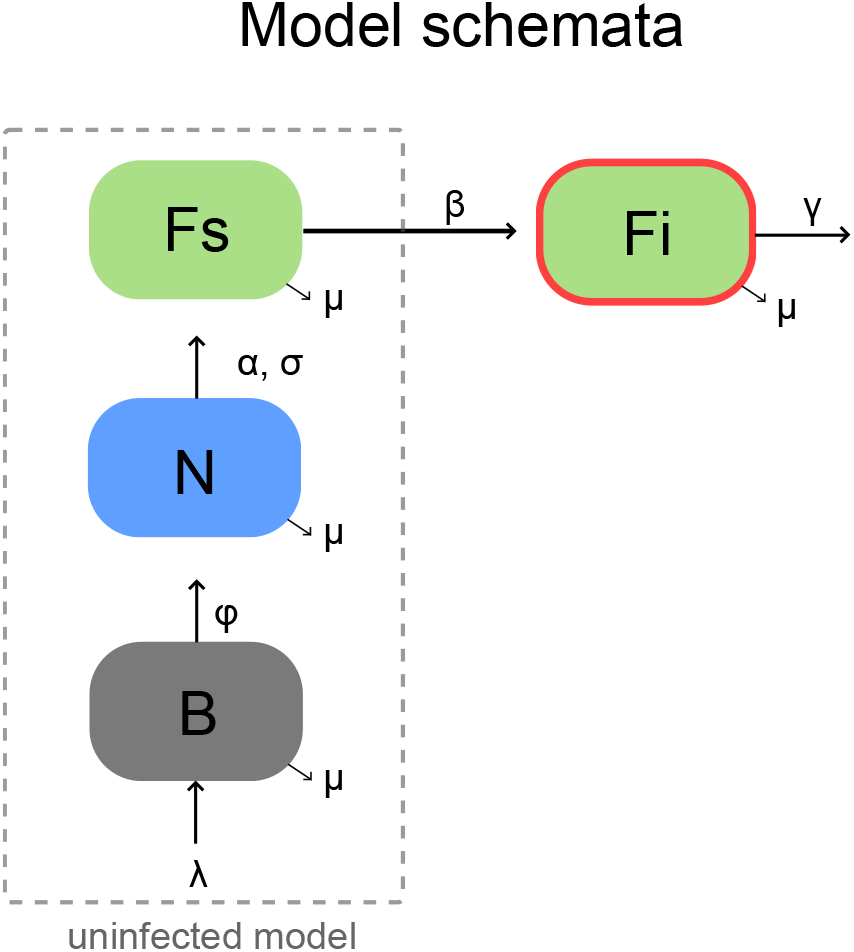
Schema for the model of colony growth in the absence and presence of *ex-nido parasitism*. Model compartments and flow. Brood (B) are born into the colony according to birth rate *λ*. Brood transition to become nest workers (N) according to the developmental rate, *φ*. Nest workers transition to become susceptible foragers (*Fs*) according to colony need (*κ*) by rate *α*. Finally, susceptible foragers can become infected (*Fi*) according to the parasite force of infection term, *β*. All compartments experience the same natural mortality rate, *µ*.

New ants are born into the brood compartment B with colony birth rate *λ* via a Hill function (see Eq. 1), which we assume is dependent on the total colony size and minimum colony size *σ* needed to maintain colony viability. We choose a Hill function to make the number of brood born into the nest conditional on the total colony size, because workers support brood care by, for example, gathering food resources, feeding larvae, and cleaning brood (10). Thus, colonies with more workers can support greater numbers of brood. Brood, B, can become nest workers, N, after a developmental period (transitioning at per-capita rate *φ*) or can die due to natural mortality (at per-capita rate *µ*). We explore a range of colony birthrates to assess the impact of the *λ* parameter on colony dynamics under the presence of *ex-nido* parasites (Fig. 4).

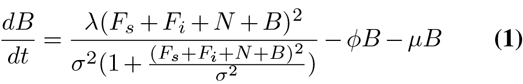

Nest workers can either die due to natural mortality at percapita rate *µ* or transition into the forager compartment, Fs, according to colony need. Here we make the assumption that a fixed proportion, *κ* of the colony is needed for extranidal tasks. We model the nest worker to forager transition using a logistic growth-like term, multiplied by the transition rate *α*, which gives the maximum per-capita transition rate (see Eqs. (2-3)). We explore how sensitive colony population dynamics are to changing this forager proportion *κ* under the presence of *ex-nido* parasitism in Fig. 5. We also perform sensitivity analyses for predicted colony dynamics under changing *α* in Fig. S4.

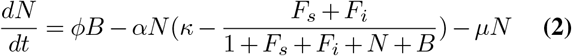

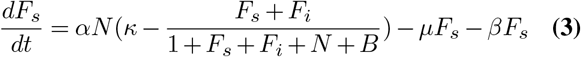

Once nest workers become foragers, we make the simplifying assumption that they remain foragers until death, which serves to reduce model complexity. For many ant species, we do not know exactly how workers are allocated to the task of foraging and how much task fidelity these workers exhibit. It is generally assumed that social insect colonies have some degree of temporal polyethism, in which workers age through a sequence of behavioral castes (10, 38) with foraging usually occurring later in life. How workers cycle through tasks over their lifetime likely depends on their physical caste as well (38, 39). While temporal polyethism gives a general pattern to how individuals cycle through tasks during their lifetime, behavioral flexibility allows colonies to be resilient during catastrophes (40–42), further complicating how colonies are socially organized. For modeling simplicity, we make the assumption that workers follow the traditional series of tasks over their lifetime (interior tasks as nest workers, followed by exterior tasks as foragers) and that foragers remain in that compartment for life. We leave investigation of the impact of behavioral flexibility observed in some colonies on interaction with *ex-nido* parasites for future work.

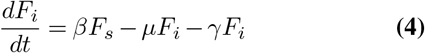

In the presence of parasitism (*β* > 0), susceptible foragers F_*s*_ become infected by parasites in the environment at rate *β*, to become infected foragers F_*i*_ (see Eq. 4). The percapita rate *β* is a parameter that combines the frequency with which foragers leave the nest, their probability of encountering parasites whilst outside the nest, and the probability of successfully becoming infected given an encounter with a parasite. Like their uninfected counterparts, infected foragers F_*i*_ can die due to natural mortality *µ*; however, they will more likely die a parasite-induced death at rate *γ*, which occurs at a more rapid rate (see Table 1 for model parameters). Though these infected foragers die at a more rapid rate than their uninfected counterparts, they do not pass on the infection, and in other respects interact with the colony like uninfected foragers. Note that infected foragers are included in the calculation of the total colony size and we assume that there are no infection-related changes in individual or colony-level behavior. In addition, infected foragers are always assumed to die (there is no recovery from infection), which is realistic given that most parasites known to infect ants require ant death as developmental necessity (13). For most results we assume that the parasite is constantly present in the environment, though we also explore the potential impacts of seasonality in the force of infection term *β*, which may arise due to seasonal fluctuations in parasite population size or prevalence. We also formally explore the impacts of changing the parasite-induced mortality rate in Fig. S3.

**Table 1.**
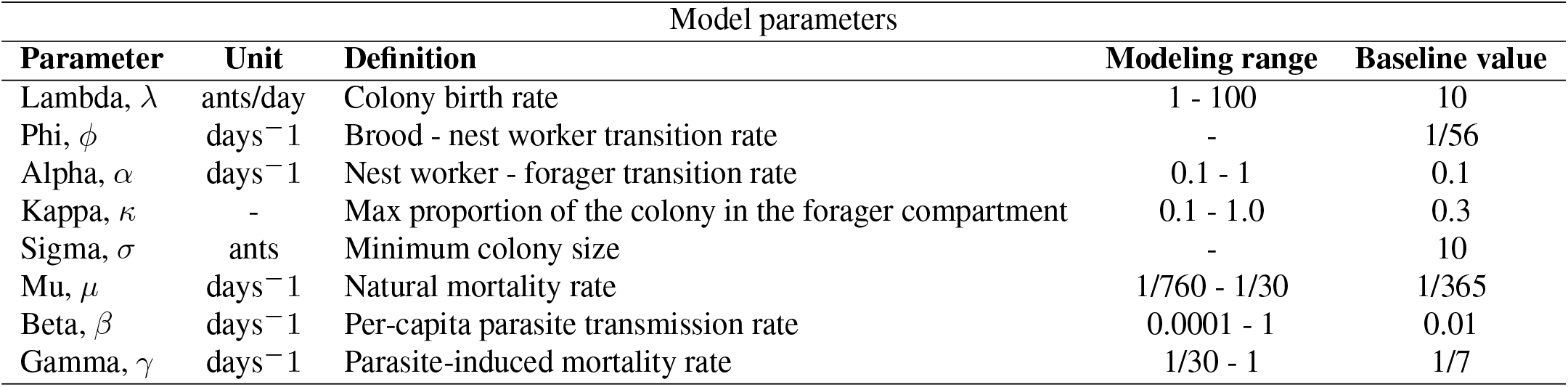
Description of model parameters, the range of their values, and the baseline values used in the modeling analysis.

### B. Additional model assumptions

#### Compartments experience identical natural mortality rates

We assume that ants in all compartments experience an identical rate of natural mortality *µ*. Empirical estimates of worker mortality rates are scarce in the literature (Table S4). While we would expect younger ants (brood and nest workers) to have lower natural mortality than foragers, due to their comparative youth and performance of less-risky tasks within the nest, that expectation has not yet been confirmed empirically, nor have the relative mortality rates been quantified. Therefore, we have made the simplifying assumption that all ants have the same lifespan of one year, corresponding to a natural mortality rate of 1/365 days^−1^. We explore the effects of changing the natural mortality rate in Fig. S2.

#### No reproductive castes are included

We do not include any reproductive castes (unmated males (drones), unmated females (gynes), or mated females (queens)) as explicit compartments in our model of mature ant colony dynamics. Reproductives do not follow the same demographic flow (i.e. brood to nest worker to forager) as non-reproductive workers and do not participate in either intranidal or extranidal working tasks (10). Furthermore, gynes and drones are expected losses for the colony; they leave on their mating flight and do not return, so whether they become infected once they have left the nest is inconsequential for colony survival (although this does have important consequences for colony fitness). The queen is implicitly included in our model, as she is responsible for the birth of new ants into the brood compartment. However, as she is a singular individual (or a few individuals, in the case of polygynous colonies), we do not create a separate compartment for her.

#### No seasonality in colony birth rate

The colony birth rate *λ* is assumed to depend on the total colony population for which we take into account all model compartments including brood and infected foragers (F_*s*_ +*N* +*B, F_s_* +*F_i_* +*N* +*B* for *β* = 0 and *β >* 0 cases, respectively), but do not take seasonal fluctuations into consideration. Many mature ant colonies fluctuate in their colony sizes and birth rate seasonally (10). Most of this seasonal fluctuation is due to the seasonal production of reproductive forms; these fluctuations are species-specific and correspond to the synchronous production and release of reproductives for mating flights between colonies. Since we do not include reproductive forms in our model, we do not include seasonality in our birth rate parameter *λ*, which also has the benefit of rendering our model more analytically tractable.

#### Only foragers are exposed to ex-nido parasites

We are investigating parasites infecting ant colonies via an *ex-nido* mode of transmission, which requires extranidal parasite exposure in order to become infected (13). In many ant societies, this means that only foragers are at risk because only foragers leave the relatively protected confines of the nest. However, some exceptions to this do occur. For nomadic species such as army ants, periodic relocation makes all ants, including brood and reproductives, susceptible to infection, particularly from parasitoids, which can conduct aerial attacks on foraging ants (43). In other cases, parasites themselves are able to enter the nest independently of the behavior of ant workers, either through mobile stages actively entering the nest (e.g. Syrphid fly microdon larvae) or through parasitoids directly ovipositing their young inside the nest (e.g. some Phorid fly species). For a first attempt at understanding how *ex-nido* parasites impact ant colony dynamics, we have chosen to only focus on the case where foragers are exposed to parasites. In future work, we hope to address the other ways that *ex-nido* transmitting parasites impact ant colony growth.

#### No seasonality in the parasite force of infection, β

For modeling simplicity, we assume that the parasite force of infection term *β* is constant, though empirical evidence suggests that parasites infecting ants are heterogeneous in both time and space (44, 45). To assess whether seasonality in the parasite force of infection term might improve colony rescue role, we further investigated how one or two annual peaks in *β* could impact infection dynamics. To do this, we use standard approaches (e.g. like that used in (46)) and assume that the force of infection term *β* takes the form *β*(*t*) = *β*_0_(1 + *β*_1_ sin(2*πt*)) or *β*(*t*) = *β*_0_(1 + *β*_1_ sin(4*πt*)), where t is in years but is adjusted to match our time scale, to get one or two annual peaks, respectively. We explore the impacts of seasonality in Figs. 6, S5, and S6.

#### Model parameters

A description of the model parameters and their values are given in Table 1. Whenever possible, estimates of model parameters were extracted from published values. We provide a detailed discussion of how model parameter values were chosen and summarize empirical estimates from the literature in the Supplemental Information and the tables therein.

## Results

We are ultimately interested in the impact of *ex-nido* parasitism on ant colony dynamics under biologically realistic parameter space. To that end, here we first provide validation of our model in the absence of parasitism, and then show how our model predicts parasite-induced impacts on colony dynamics under changing values of colony birth rate and parasite force of infection. Finally, we assess model sensitivity to different parameter values such as changing the proportion of the colony foraging, changing the natural mortality rate, and adding seasonality to the parasite force of infection.

### C. Baseline model validation

To understand the effect of *ex-nido* parasitism on ant colony dynamics we first examined dynamics in the absence of parasitism (force of infection *β* = 0 days^−1^), using baseline parameter values given in Table 1. Using linear stability analysis and confirmation via direct numerical simulation, we found that for all birth rates in the range of modeled values, colony growth reaches equilibrium in approximately 1,500 days, which corresponds well to knowledge of ant colony development from inception to colony maturity (8, 10, 32, 47, 48). To assess how the model reacts to population perturbations in the absence of *ex-nido* parasitism, we simulated the removal of a fixed percentage of each compartment (B, N, F_*s*_) after equilibrium had been reached (Fig. S1). In all cases, as long as the population had not been reduced below the minimum value necessary to sustain colony functioning (*σ*, Table 1), each compartment was able to recover back to their equilibrium values. The time it took to recover from the perturbation to 90% of equilibrium values was 600 - 820 days (2 - 3 years) and 1,110 - 1,650 days to return to full equilibrium values. While it is unknown how long it would take an ant colony to recover back to pre-perturbation population values in a natural setting, it seems realistic that the recovery period is on par with the growth period of an incipient ant colony reaching maturity.

### D. Equilibrium values and bifurcation diagrams

The equilibrium values for each compartment within the uninfected model (*β* = 0 days^−1^) under baseline parameter values (Table 1) are as follows-Fs: 1,045.91 ants, N: 2,118.56 ants, and B: 485.507 ants, which is a realistic colony size for many ant species (7, 10). The equilibrium values for each compartment within the infected model (*β* = 0.01 days^−1^) under baseline parameter values (Table 1) are F_*s*_: 458.134 ants, F_*i*_: 31.4659 ants, N: 1,034.09 ants, and B: 485.499 ants. Bifurcation diagrams for the model when *β* = 0 (uninfected) and when *β >* 0 (infected) are provided in Figs. 3a and 3b, respectively. In Fig. 3a, the bifurcation diagram switches from a stable to unstable regime at the minimum colony size needed for functioning (min = 10 ants). In Fig. 3b, the bifurcation diagram with respect to *β* remains stable throughout the range of *β* values for a colony birth rate *λ* = 10 ants/day.

**Fig. 3.**
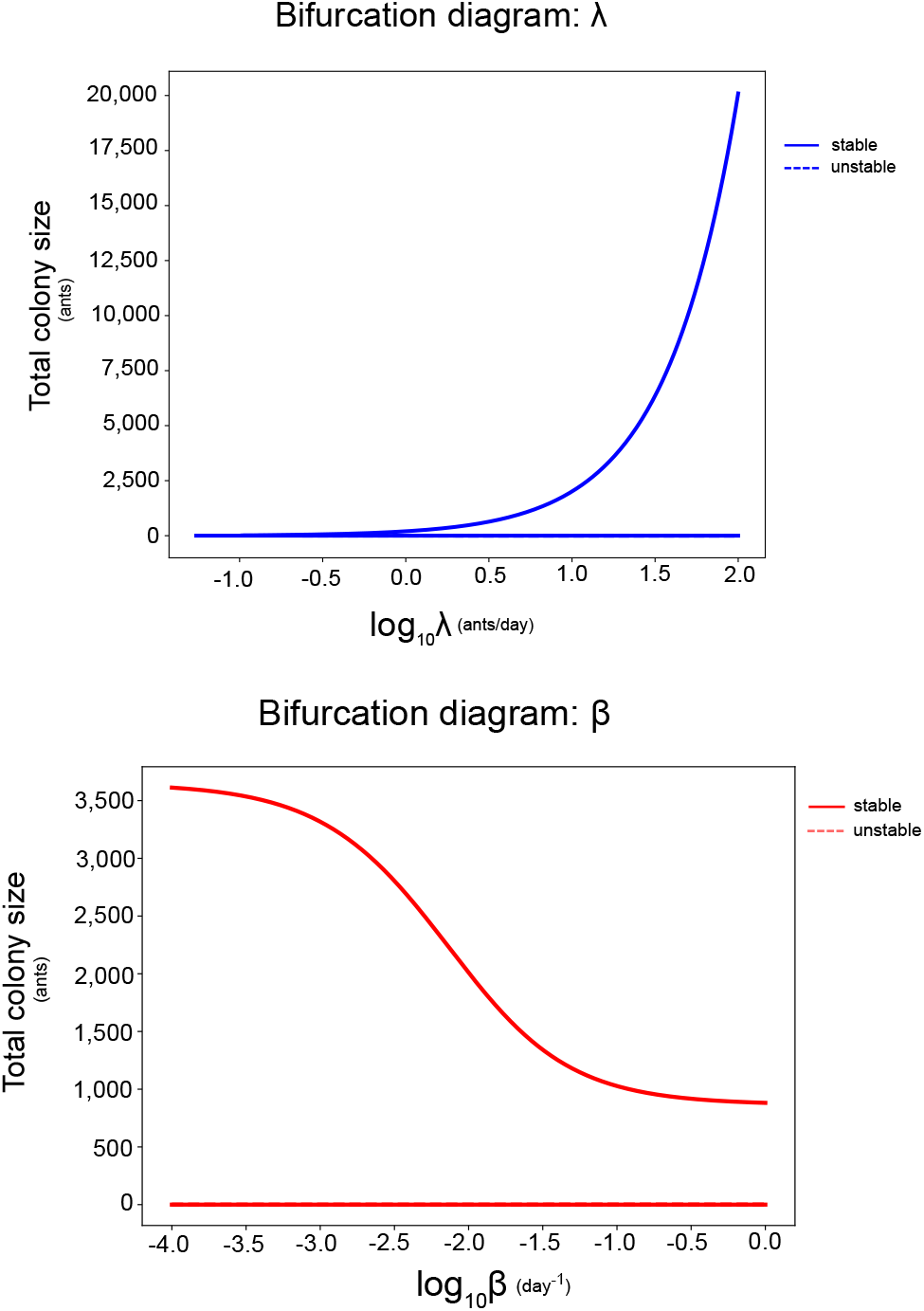
Bifurcation diagrams with respect to colony birth rate *λ* and parasite force of infection, *β*. (a) Bifurcation diagram for the uninfected model (eqs. 1-3, *β* = 0) under baseline parameter values given in Table 1. Rescaled colony size as a function of colony birth rate. Colony size was rescaled using an arcsinh function. (b) Bifurcation diagram for the infected model (eqs. 1-4) under baseline parameter values given in Table 1.

### E. Changing birth rate, *λ*, and force of infection, *β*

We next examined colony dynamics in the presence of *ex-nido* parasitism. Under the baseline colony birth rate and force of infection given in Table 1, we observe an equilibrium reduction of 45.8% in colony size in the infected model relative to the uninfected model (i.e. *β* = 0 days^−1^). It takes approximately 2,900 days to reach this full equilibrium reduction and approximately 800 days to be within 10% of equilibrium values. When *β* is fixed at the baseline value of *β* = 0.01 days^−1^ and we assume different values of colony birth rate *λ* from 1 - 100 brood/day, we predict that the proportion reduction in colony size remains essentially the same (Fig. 4a), and it takes 700 - 900 days to be within 10% of equilibrium values. When we fix colony birth rate *λ* at the baseline value of *λ* = 10 brood/day, chosen because it’s in the middle of log scale realistic values for colony birth rates (see Table S1) and traverse values of *β* in the range of *β* = 0.0001 to 1 day^−1^, we find a wide range in resulting proportion reductions in colony size, from 1% up to 77%, respectively (Fig. 4a). This is realistic because *β* controls the rate at which foragers are lost due to parasitism; above a certain value of *β* the colony’s birth rate can no longer compensate for the loss of foragers and thus larger reductions in colony size occur (Fig. 4).

**Fig. 4.**
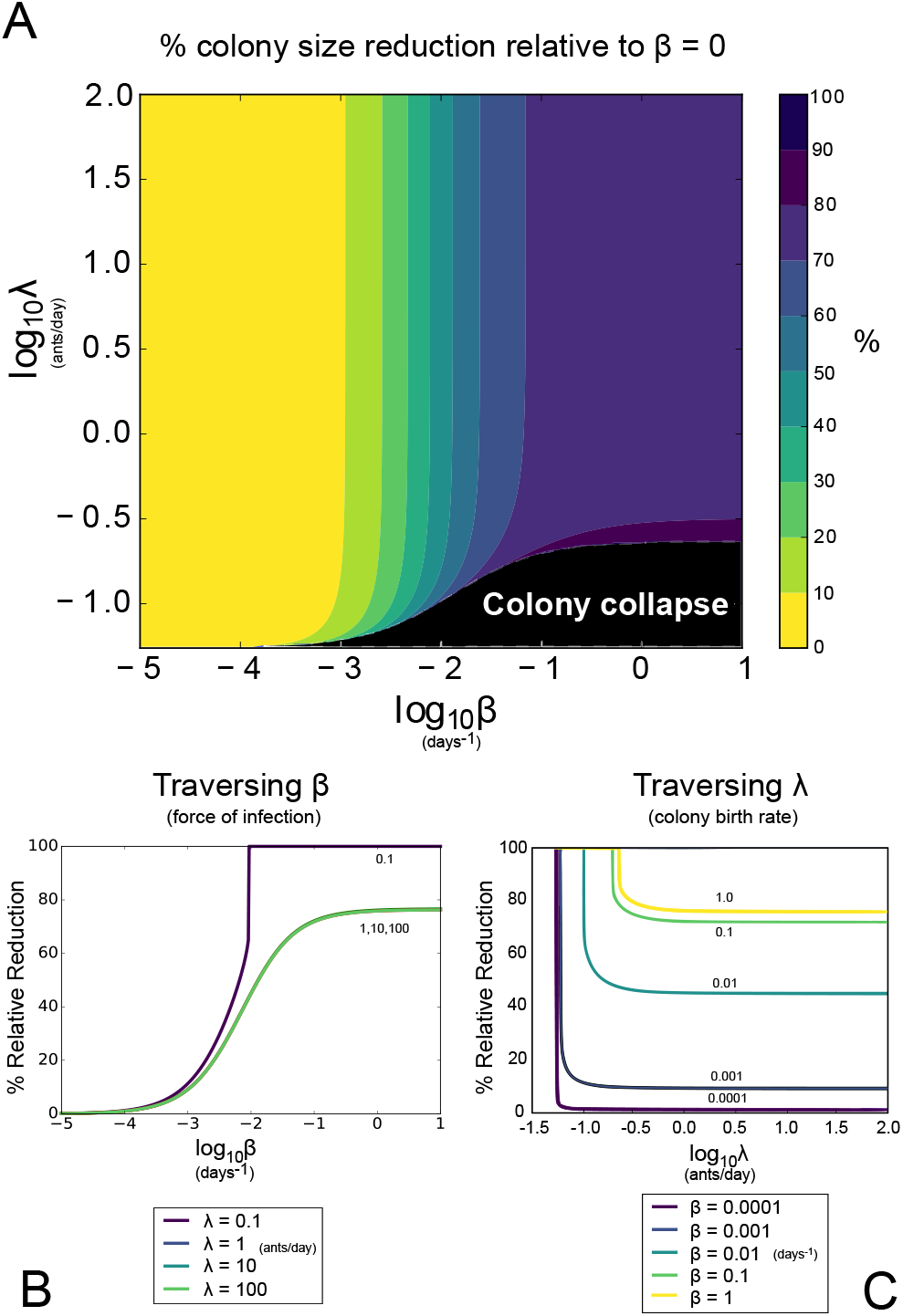
Predicted impact of *ex-nido* parasitism on colony growth dynamics. (a) Contours in percent reduction in colony size predicted by our model (Eqs. (1-4)) as a function of colony birth rate *λ* and force of infection *β* relative to when *β* = 0. Other parameters can be found under baseline values in Table 1.(b) Percent reduction in colony size predicted by our model (Eqs. (1-4)) as a function of rate of infection *β* and *λ* = 10 brood/day relative to when *β* = 0. Other parameters can be found under baseline values in Table 1.(c) Percent reduction in colony size predicted by our model (Eqs. (1-4)) as a function of colony birth rate *λ* and *β* = 0.01 relative to when *β* = 0. Other parameters can be found under baseline values in Table 1.

### F. Impact of the proportion of the colony foraging, *κ*

The proportion of the colony that forages *κ* is an important model parameter because it impacts the number of individuals in the susceptible forager F_*s*_ compartment and thus the number of individuals at risk of becoming infected. Altering the proportion of the colony foraging *κ* had a large impact on the resulting percent reduction in colony size relative to the uninfected model (Fig. 5a) and on the total colony size at equilibrium (Fig. 5b). When 10% of the colony forages under baseline model conditions, there is a 22.8% reduction in colony size compared to the uninfected model; in contrast, when 70% of the colony forages, there is a 61.3% reduction in total colony size relative to the uninfected model (Fig. 5a). This translates into a large difference in resulting total colony sizes for a typical ant colony: 2,816 ants vs. 1,412 ants, for *κ* = 0.10 vs. 0.70, respectively. By allowing a larger proportion of the colony to be in the susceptible forager (F_*s*_) compartment, more individuals are at risk for becoming infected, and so a larger number of individuals die due to parasite-induced mortality rather than natural mortality. Our model, which makes the simplifying assumption that foragers experience the same natural mortality as their non-foraging counterparts, likely overestimates the percent relative reduction in colony size between infected and uninfected models because foragers likely do experience a higher rate of natural mortality.

**Fig. 5.**
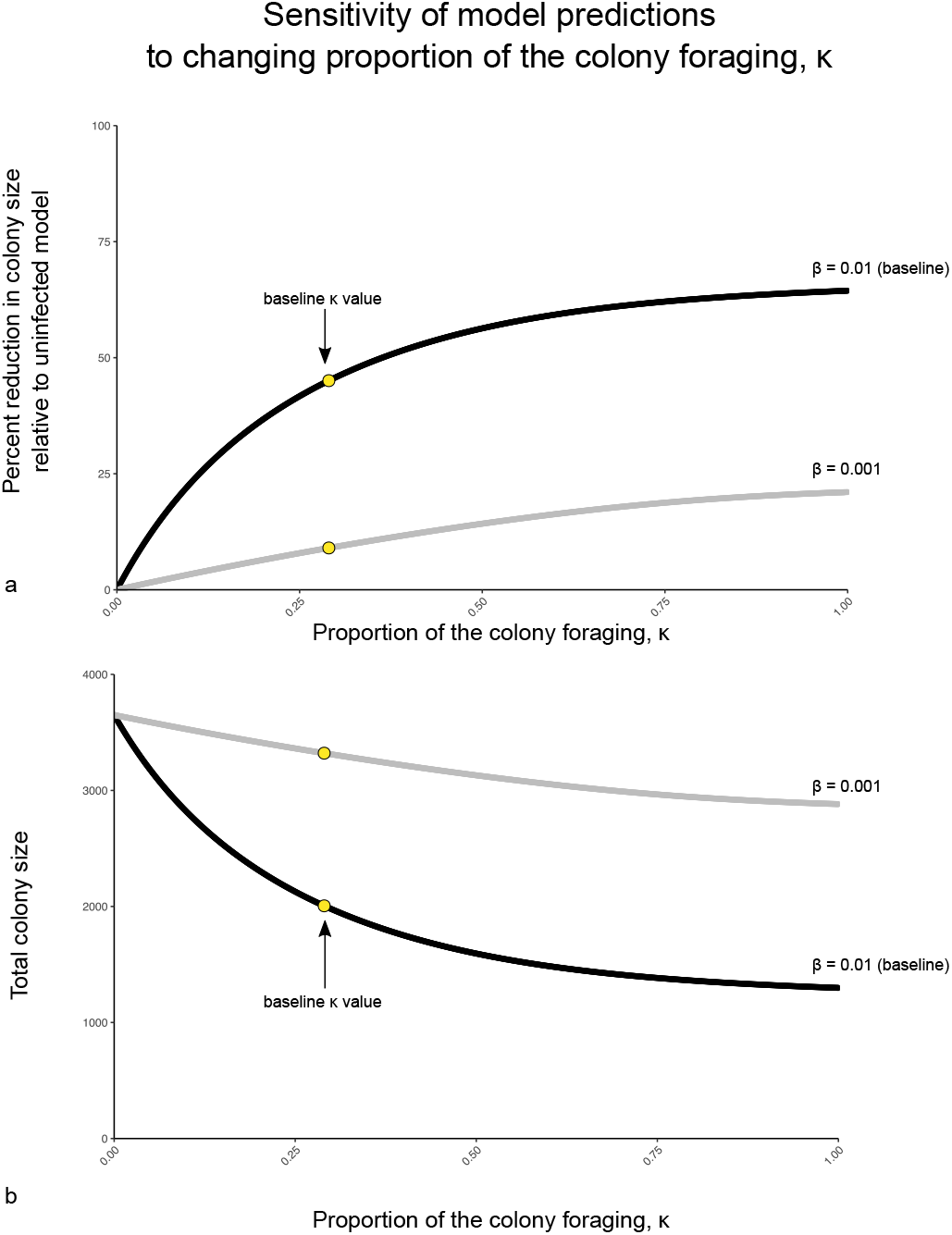
Sensitivity of model predictions to changing the proportion of the colony foraging, *κ*. Model predictions are from numerical simulation under baseline parameter values given in Table 1. (a) The proportion reduction in colony size relative to the uninfected model under changing values of *κ* from 0 to 1. (b) Total colony size (Fs + N + B) under changing values of *κ* from 0 to 1.

### G. Adding seasonality to parasite transmission rate

To assess how seasonality in the parasite force of infection term *β* could impact colony dynamics, we simulated outbreaks in which *β* varies over time rather than being held at a fixed value. To accomplish this, in these simulations the force of infection term *β* takes the form *β*(*t*) = *β*_0_(1 +*β*_1_ sin(2*πt*)) or *β*(*t*) = *β*_0_(1 + *β*_1_ sin(4*πt*)), to get one or two seasonal peaks per year, respectively. We plot these seasonally varying values of *β* and their impacts on the percent colony reduction relative to the uninfected model in Figs. S5 and S6 and we show a comparison of model predictions for total colony size in cases where *β* is either constant, has 1 seasonal peak per year, or has two seasonal peaks per year in Fig. 6. When *β* reaches a seasonal peak, colony sizes in all three cases approximately mirror each other. However, including seasonality in the force of infection term *β* allows colony size to rebound slightly, resulting in a higher average colony size over time Fig. 6. Thus, our results in Fig. 4a, in which parasitism is assumed to be constant, represent the ‘worst-case’ scenario for a given value of *β*.

**Fig. 6.**
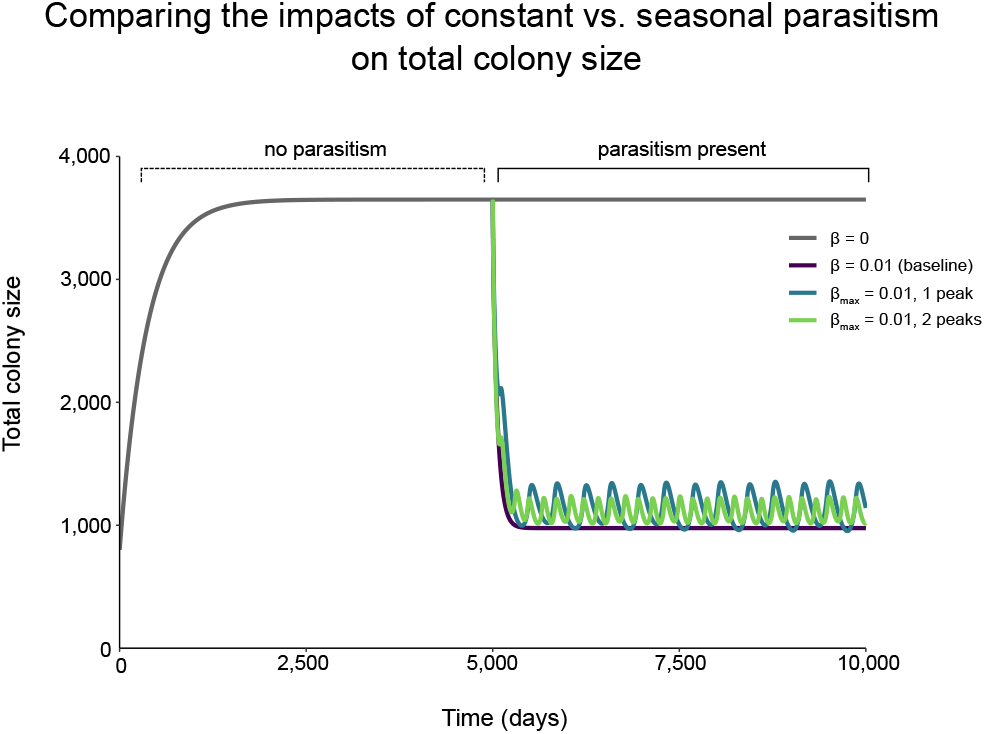
Comparing the impact of constant vs. seasonal parasitism. (a) Values of the force of infection term *β* under constant and seasonal parasitism with one or two peaks. The maximum values of *β* = 0.19 days^−1^. (b) Model predictions of colony size when there is no parasitism present, parasitism is present under baseline values that are constant, parasitism fluctuates seasonally with one annual peak, or parasitism fluctuates seasonally with two annual peaks. Model predictions are from numerical simulation under baseline parameter values given in Table 1 and *β* = 0.19 days^−1^.

### H. Changing the natural mortality rate *µ*

As the natural mortality rate decreases (i.e. individuals live longer), the total population increases markedly (Fig. S2b). When *ex-nido* parasites are present (*β >* 0), changing the natural mortality rate can also cause large differences in the percent reduction in total colony size between the infected and uninfected models (Fig. S2a). For example, when we take an average ant lifespan of 1 year (*µ* = 1/365 days^−1^), which has been reported for worker ants, the percent relative reduction in total colony size is approximately 44%, whereas when the average lifespan is assumed to be 30.4 days (*µ* = 1/30.4 days^−1^), the percent relative reduction is approximately 3%. An average longer lifespan increases the relative impact of parasitism on the colony in two ways. Firstly, foragers live longer and thus have more cumulative exposure to parasites over the course of their longer lifespan, resulting in a larger number of infected foragers at equilibrium. Secondly, as the lifespan of individuals increases, there is a larger disparity between the parasite-induced mortality rate *γ* and the natural mortality rate *µ*, resulting in a larger disparity when comparing the colony population sizes of the infected and uninfected models. In the uninfected model, fewer individuals are dying due to natural mortality, so there is a larger population size, while population loss in the infected model is being driven primarily by parasite-induced mortality.

### I. Changing parasite-induced mortality rate, *γ*

Changing *γ*, the parasite-induced mortality rate for infected foragers, alters the equilibrium number of infected foragers. In our model, infected foragers are assumed to behave as healthy foragers, don’t infect others, and are considered part of the total population size that is used to calculate the fraction of the colony that is foraging. Thus, when the parasite-induced mortality *γ* is lower (but still faster than the rate of death due to natural mortality), infected foragers survive longer and replacements do not need to be recruited from the nest worker population as frequently, which helps reduce the size of the forager population that is exposed to infection at rate *β*. However, this difference has only a small impact on the percent reduction in colony size relative to the uninfected model (Fig. S3a) and the total colony size at equilibrium (Fig. S3b). When *γ* is 1/30 days^−1^, the resulting percent population reduction is 43.3%, while the percent reduction only increases slightly to 46.6% when *γ* is changed to 1 days^−1^ (Fig. S3a). Thus, model outcomes are not very sensitive to changes in *γ* within biologically reasonable ranges.

### J. Conditions for colony collapse

Given specific colony parameters, we can use our model to predict colony size reduction or collapse. In the case of the baseline parameters (Table 1), colonies are vulnerable to collapse when the birth rate *λ* is less than 0.2328 ants/day, which we calculate via linear stability analysis of Eqs. (1 - 4) in the limiting case of *β → ∞*. That is, if *λ* is above this threshold then collapse will not occur, though the extent of colony size reduction will depend on the strength of the force of infection *β* (Fig. 4a,b). In Fig. 4b, the only value of *λ* that results in collapse is less than this threshold, and in Fig. 4a, the upper boundary of the collapse region asymptotically approaches this value. If *λ* is less than the threshold value of 0.2328 ants/day, the colony may collapse if *β* is sufficiently large. Though the total colony size at equilibrium is sensitive to changing values of *κ*, the birth rate threshold above which collapse will not occur is not sensitive to changing values of *κ*. For example, for *κ* = 0.01, the minimum value of *λ* is 0.0656 brood/day, whereas for *κ* = 0.5, the minimum value of *λ* to avoid collapse is 0.2736 ants/day. While ant colony birth rates are species-specific and likely do vary seasonally, a birth rate of 0.2328 ants/day is much lower than those reported in the literature (Table S1). For much of the parameter space explored in our model (Fig. 4a), colony collapse is not predicted to occur.

## Discussion

The ants and other social insects are complex societies that have evolved over long periods of evolutionary time (6). For the ants we know this to be between 139 - 158 million years, which parallels major changes in ecological complexity, such as the radiation of the angiosperms and expansion of other insect groups (49). During this time period, ants evolved to be the dominant animals in most terrestrial biomes; despite accounting for 0.4% of the estimated 5.5 million insect species (50), they typically comprise *>* 50% of animal biomass (15, 51). This success seems paradoxical, as ants are host to diverse parasites (12, 13, 33) and seem to be particularly vulnerable to infectious disease outbreaks due to living in crowded conditions with lots of highly related individuals. Accordingly, a lot of research has been devoted to uncovering potential mechanisms by which ants and other social insects might mitigate potential disease spread inside their colonies (reviewed in (12, 16, 18, 19)). However, the majority of the parasites that infect ants and social insects use lifecycles that preclude intra-colony transmission, instead requiring periods of time outside the nest for development in the environment or in other hosts before the next transmission event to ants can occur (12, 13). The impact of these parasites on colony dynamics has been hitherto unexplored.

Here we formally explored the potential for epidemic collapse due to these *ex-nido* parasites by modeling the impact of their infection on ant colony dynamics. Using parameter values extracted from the literature (Appendix B), we found that in the absence of parasitism, colonies can recover from population perturbations back to equilibrium values within approximately 1,100 - 1,650 days (Fig. **??**), which is in general accord with reported growth rates of colonies as they mature (47). Such perturbations might reflect attack by predators or competition with other colonies. We then modeled the presence of *ex-nido* parasites and showed that for a wide range of biologically plausible birth rates the effect of parasitism ranged from negligible impacts on colony population (*<* 10% colony size reduction, Fig. 3a) to significant reductions (>90% colony size reduction, Fig. 3a), depending on the magnitude of the force of infection parameter *β*. Under the baseline parameters used in our model, colonies are not vulnerable to parasite-induced collapse unless *λ* is below 0.2328 ants/day, which is much lower than birth rates reported in the literature (Table S1). Thus our modeling predicts that under realistic parameter values, infection by *ex-nido* parasites will not result in colony collapse.

An important question is whether values of *β* large enough to cause collapse could be achieved. A key result of this study is that the impact of *ex-nido* parasitism depends most heavily on the force of infection *β*, a parameter that combines the frequency with which foragers are in the extranidal environment, how frequently foragers encounter parasites while in that environment, and the probability of successful infection given contact with a parasite. While few if any empirical estimates of the components of *β* are known (see additional discussion in SI), if foragers make trips very often, encounter parasites very frequently, and/or are unable to prevent successful infection by behavioral, social, or physiological immune mechanisms, then such values could indeed occur. The little information we do have on the individual components of *β* suggests that the parasite infective pressure that ants face is probably quite heterogeneous in both time and space (52–54). For modeling simplicity we have assumed, for our main results, that parasite pressure is constant, but many parasites of social insects undergo seasonal fluctuations in their populations (45, 55, 56). When we investigated seasonality by allowing for one or two annual peaks in parasite force of infection *β*, colonies were able to rebound and colony size were slightly larger on average (Fig. 6). Our model, which assumes a constant rate of parasite infection, therefore represents an upper bound to the impact of *ex-nido* parasite pressure on ant colony size. Additionally, we know that ants are able to behaviorally mitigate their potential exposure to parasites by reducing foraging (57–59) and can prevent exposure from developing into infection via self- and allogrooming behaviors (25, 60). Our model, which does not include behavioral flexibility or behavioral anti-parasite defenses, likely overestimates the impact of parasitism. Thus, the conditions that lead to model-predicted colony collapse provide a worst-case scenario for the colony, and these conditions are unlikely to occur in natural settings.

Furthermore, the timing over which these parasite-induced colony size reductions occur plays an important role in colony robustness; these reductions take place over years, not in a matter of days (approximately 800 days or over 2 years to be within 10% of full equilibrium reduction under baseline parameters), causing a gradual thinning of colony size. This contrasts with what we observe in high density, group living mammals where a sudden catastrophic loss of individuals can occur, like that recently observed during the Saiga antelope epizootic (61) or occurring in tropical forest mammal populations due to anthrax (62). In natural settings, ant colonies might be able to adapt to or mitigate this gradual parasite-induced loss of workers before collapse occurs, through changing behavioral defenses (57), moving nest locations (63), or potentially increasing birth rate in compensation.

What happens in cases where colony size is reduced by parasitism but collapse does not occur? Few studies have explored the impact of parasite-induced colony size reduction on colony functioning, organization, or fitness. Ants and other social insect colonies do have built-in redundancy to buffer loss (10, 32, 64). Schmid-Hempel and Heeb (65) found that imposing an extra 10-15% weekly mortality rate on *Bombus lucorum* colonies did not alter colony growth rate or reproductive output compared to control colonies. Followup work by Müller and Schmid-Hempel (66) experimentally removed 10% of the worker force each day in *B. lucorum* colonies and found no difference in worker production or the timing of reproduction but there were reductions in either the number of males produced or female reproductive body size depending on when worker loss occurred. One proxy for forager loss may be circumstances where foragers do not leave the colony for resources because of risk of death from abiotic factors like heat. In a long term study on the desert foraging ant *Pogonomyrmex barbatus*, Gordon (67) showed that colonies could reduce the number of foraging trips to avoid worker desiccation and still achieve colony growth and colony fitness (number of daughter colonies produced).

Are there any examples of parasites causing epidemic colony collapse in natural populations? For the ants, to our knowledge there are no reports of parasite-induced colony collapse for mature colonies, but there are examples of colonies persisting despite the multi-year presence of *ex-nido* transmitting parasites (54). While attempts to use both *in-nido* and *ex-nido* transmitting parasites as biocontrol agents against pestiferous ant species have been numerous (see review by (68)), their lack of success highlights how difficult it is to perturb a mature ant colony to collapse (69–71). Parasite pressure is most certainly a significant selective pressure on incipient colonies (10), however; this is extremely hard to observe in field settings and so associated data is extremely limited.

Some social insects are famously experiencing collapse. Managed bee colonies and global bee populations have been experiencing significant decline beginning in the early 2000’s (72, 73), but it appears that a constellation of causes including pesticide usage, habitat loss, chronic stress, and directly transmitting parasites coupled with ectoparasites (mites) contributed synergistically to the reported losses (74). Our understanding of the parasite pressure that social insect colonies face in nature remains limited, but it seems that parasite-induced collapsed of mature social insect colonies in natural populations are rare events, and our model suggests that parasite-induced collapse by *ex-nido* transmitting parasites would take years, not days or months, to occur.

What makes ant and other social insect colonies so resilient in the face of possible disease threats? Unlike other animal societies and group living organisms, ants and social insect colonies employ an efficient division of labor that allows for the loss of individuals while maintaining colony functioning (10, 32, 64). Furthermore, it is this very division of labor that helps mitigate disease exposure by limiting the number of individuals who leave the protected confines of their nest, a form of ‘organizational immunity’ (18). Finally, strong physiological and social immune defenses inside the nest help prevent the potential onward transmission of directly transmitting parasites (reviewed in (12, 16–19)).

## Conclusions

Ants and other social insects have had to contend with the intense selective pressure imposed by parasites and pathogens over their long evolutionary history. The ability to prevent outbreaks likely reflects early selection prior to their transition to a eusocial lifestyle (23). This is especially important for the ants, which evolved colony life from a starting position in microbe rich soils (75), likely necessitating possessing a strong prophylactic disease defense system. Work has shown that ants and social insects do have an impressive suite of behavioral and physiological immune mechanisms at their disposal (12, 16–19). The evolution of a eusocial lifestyle, in which fitness lies at the level of the colony and not at the level of the individual, coupled with these strong anti-parasite defenses and lots of evolutionary time, has likely facilitated the co-evolution of *ex-nido* transmission strategies on the part of their parasites (76), which remains an exciting and underexplored area of research. Our work here shows that parasites that require leaving the nest before the next transmission event to ants can occur are likely to have a limited impact on mature colony dynamics, precluding epidemic collapse. This then may be a key difference between social insect societies and other animal societies, where massive outbreaks of infectious diseases are routine and destabilizing to group functioning.

## ACKNOWLEDGEMENTS

We would like to thank Timothy C. Reluga for technical guidance in modeling analysis and for insightful discussions thereof. We would also like to thank members of Penn State’s Center for Infectious Disease Dynamics for helpful comments and feedback on this work. We gratefully acknowledge the many natural historians who have collected records of parasites infecting ants as well as the myrmecologists who have studied ant life history traits. Their work provides the biological foundation on which this and other studies can be built.

This material is based upon work supported by the National Science Foundation (NSF) under Grant No. 1414296 to DPH as part of the joint NSF-NIH-USDA Ecology and Evolution of Infectious Diseases program and under NSF Grant No. DMS-1714654 to JMC. LEQ is supported by a NSF Graduate Research Fellowship under Grant No. DGE1255832. Any opinions, findings, and conclusions or recommendations expressed in this material are those of the authors and do not necessarily reflect the views of the National Science Foundation.

## Supplementary Note 1: Model Parameters

A description of the model parameters and their values are given in Table 1. Whenever possible, estimates of model parameters were extracted from published values. Here we provide a detailed discussion of how model parameter values were chosen, and summarize empirical estimates from the literature in Tables S1, S2, S3, S4, S5, S6, and S7.

### Colony birth rate *λ*

For colony birth rate, there is a large range in published values in the literature. This is due to large differences between ant species, with some species having extremely large colony sizes and thus large numbers of eggs laid per day while other species have small colonies and smaller birth rates (Table S1). In our model, the overall rate at which new brood are born into the colony is dependent on the colony birth rate parameter *λ*, the current colony population (Fs + Fi + N + B), the minimum colony size *σ* needed to maintain colony viability, and this together is modulated via a Hill function (see Eq. 3.1). We explore a range of colony birthrates to assess the impact of the *λ* parameter on colony dynamics under the presence of *ex-nido* parasites (Fig. 4).

### Brood - nest worker transition rate *ϕ*

For the brood-nest worker transition rate, *ϕ*, the range in this parameter is due to the impacts of temperature on brood development time, with values of 18 days to 180 days reported in the literature (Tables S2, S2b). For simplicity, we assume a constant transition rate with value 1/56 days^−1^, the inverse of the median brood development times reported in the literature.

### Nest worker - forager transition *α* and *κ*

The nest worker-forager transition is governed by two parameters in our model: *κ* (the maximum proportion of the colony that are foragers), which dictates how many new foragers need to be recruited from the nest worker compartment, and *α*, the maximal rate at which the nest worker-forager transition occurs in the absence of foragers. Estimates of the proportion of ant colony populations that engage in foraging are scarce (Table S3), and the estimates that have been reported vary considerably, ranging between 4.5 - 90%. It is likely that the proportion of the colony that foragers changes seasonally, but for model simplicity we make the assumption that a maximum of thirty percent of the colony is in the forager compartment at any given time. We explore how sensitive colony population dynamics are to changing this forager proportion *κ* under the presence of *ex-nido* parasitism in Fig. 5. We also perform sensitivity analyses for predicted colony dynamics under changing *α* in Fig. S4.

### Natural mortality rate *µ*

Our understanding of natural mortality rates and worker lifespan in ants is limited by the relative difficulty of following individuals over the course of their lifetime under field settings. Accordingly, most estimates of worker lifespan come from laboratory studies, which preclude many natural causes of death (i.e. predation, interspecific competition and aggression, and parasitism). Worker life spans greater than one year have been reported for several species (Table S4), with some exceeding three years. For studies that have followed worker life spans in the field, the life spans of ants once they become foragers are far less (6 - 30 days). For simplicity, we assume an individual’s lifespan to be 1 year, which corresponds to a natural mortality rate of 1/365 days^−1^. We assume that each compartment has the same natural mortality rate; we explore the effects of changing the natural mortality rate in Fig. S2.

### Parasite-induced mortality rate *γ*

In our model, we assume that successful infection leads to mortality, thus we define parasite-induced mortality rate as the rate at which infected individuals die, not their probability of dying. Few estimates of parasite-induced mortality rates for parasites of social insects exist in the literature. We summarize reported estimates of the inverse survival time in Table S5 and formally explore the impacts of changing the parasite-induced mortality rate in Fig. S3.

### Supplementary Note 2: Additional modeling results

#### Changing the nest worker - forager transition term *α*

Changing the nest-forager transition term *α* does not qualitatively change either the transient or equilibrium dynamics for the model (Fig. S4). An *α* of 0.1 leads to a percent reduction of approximately 46% relative to the uninfected model, whereas an *α* of 1.0 leads to a percent reduction of approximately 50% (Fig. S4), so the model is not very sensitive to changes in *α* within biologically reasonable ranges. Altering *α* slightly modifies how quickly nest workers transition into the forager compartment depending on colony need (i.e. to make up the difference between the number of foragers present and the foraging cap *κ*). Smaller *α* indicates that the transition to fill the forager compartment occurs more slowly; this results in slightly fewer foragers being in the compartment leading to fewer individuals being at risk of becoming parasitized and thus a slightly reduced percent reduction in total colony size.

### Supplementary Note 3: Additional Discussion

Please refer to the main text for a complete discussion of the results of this study. Here we present additional discussion fodder for interested readers.

#### The force of infection parameter *β*

The results of our model show that the force of infection parameter *β* is extremely important for predicting the severity of the impact of indirectly transmitting parasites on total colony size. The *β* parameter is composed of three different parameters: the rate at which individuals forage, the rate at which foragers encounter parasites in the extranidal environment while foraging, and the probability that an encounter with a parasite while foraging becomes a successful infection. For all three components of *β*, few if any empirical estimates are known. While work has been done on colony-wide foraging ranges and rates (77, 78), the frequency with which a given individual forages has been measured only a handful of times (79–82). Studies investigating individual foraging rates suggest that most foragers make few trips while a small proportion of foragers make many trips (79), which would lead to heterogeneity in infection risk. It is also important to acknowledge that non-foragers can also be periodically exposed to the extranidal environment and thus to parasite exposure. For example, in nomadic ant species such as those in the army ant tribe Ecitonini, frequent nest relocation could expose brood and intranidal workers as well. How such host ecological traits relate to parasitism pressure will be the subject of future work. The second component of *β*, how frequently parasites are encountered in the extranidal environment, remains enigmatic. For the vast majority of parasites infecting social insects, we have no idea how frequently potential hosts encounter infective stages such as fungal spores, largely due to the inherent difficulties in gathering such data. Some work has investigated the attack rates of parasitoids and found that attack rates can vary with host density and distance from nests (83) as well as with the number of host species in the area (84). For the ant fungal parasite *Ophiocordyceps unilateralis s.l*., infected cadavers have been behaviorally manipulated to die above foraging trails at the doorstep of the colony (54), where they can rain spores down on foragers as they leave the nest. However, these cadavers have been shown to have temporal windows over which spores are released and as foragers continue to walk over trails spores are effectively ‘cleaned up’, suggesting variable individual encounter rates. Thus, while information on actual encounter rates is scarce, heterogeneity appears to be likely in how frequently social insect hosts encounter parasites in the extranidal environment.

The third component of *β* - how often parasite encounters lead to successful infection - is also relatively unknown, but host behavior and the effects of social immunity could serve to significantly reduce actual parasitism rates. When parasitoid flies are present, ants have been shown to reduce their foraging behaviour in response (58, 85, 86) and leaf-cutter ants even have workers who defend foragers from parasitoid attacks (87). For individuals that have picked up fungal spores or infective larval stage helminths, allogrooming and self-grooming can be effective strategies to prevent actual infection (24, 60, 88–90). Other mechanisms of social immunity, such as the spraying of formic acid and use of metapleural gland secretions (91, 92) can also kill parasites before infection can commence.

Taken together, what information we do have on the above components of *β* suggests that the infective pressure of parasites is probably quite heterogeneous. Thus, our model, which uses a constant rate of parasite infection likely overestimates the impact of indirect parasite pressure on the reduction of colony population numbers. Better empirical estimates of the components of *β* are needed to truly identify the impact *ex-nido* parasites might have on colony populations.

#### Impacts of worker loss on colony functioning and fitness

This work has explored how the presence of indirect parasites could impact colony population dynamics, but how such potential population reductions translate into impacts on colony fitness remains an open question. Does a 10% reduction in colony population lead to a 10% reduction in colony fitness? This seems unlikely but empirical data are scant. Also, are colonies able to functionally buffer the loss of individuals through redundancies in their social organization? Indirect parasites, by modifying host behavior (i.e. reduced foraging in the presence of parasitoids (58), can also potentially impact colonies in nuanced ways that are subtler than simple reductions in colony size. Better understanding of the link between colony size, social structure, functioning, and fitness will allow us to better understand the interaction of social insects and their parasites in an evolutionary context.

The impact of worker loss might manifest in other ways besides colony growth or the production of reproductives. Increasing evidence shows that there are important individual differences in task performance and personality (93), and these differences can cascade up to alter colony-level organization and phenotypes (94, 95). Thus, the parasite-induced loss of ‘keystone’ individuals could potentially alter colony phenotype, with downstream consequences for colony competitive ability and thus growth, survival, and reproduction.

An additional way that the parasite-induced loss of workers could impact colonies is by promoting a positive feedback between worker loss and future parasitism events if, for example, inexperienced foragers had to make more trips or encountered more parasites in the environment than the experienced foragers that they replaced. While our model does not include this potential feedback, it would be interesting to examine both theoretically and empirically.

#### Impacts of parasites that cause morbidity rather than mortality

For modeling simplicity, we have only included parasite records where the parasite clearly caused mortality for the infected individual. We have excluded parasites that cause sickness behavior rather than mortality, or where the exact extent of host pathology is unclear. The colony-level impact of these parasites causing morbidity merits further investigation.

#### Need for more empirical measurements of ant colony sociometry

The modeling approach employed in this work has been useful for clarifying what host biological features might be the most important for colony survival while in the presence of indirect parasites. While some parameter estimates were easily gleaned from published studies (i.e. brood developmental time), other important parameters such as colony birth rate and the proportion of the colony work force dedicated to foraging have far less empirical data available. In order to more accurately model how colonies grow in the presence of absence of parasitism and to validate this current model, we need better estimates of basic ant colony sociometry, as has been advocated by Tschinkel (96).

### Supplementary Note 4: Supplemental Figures and Tables

**Fig. S1.**
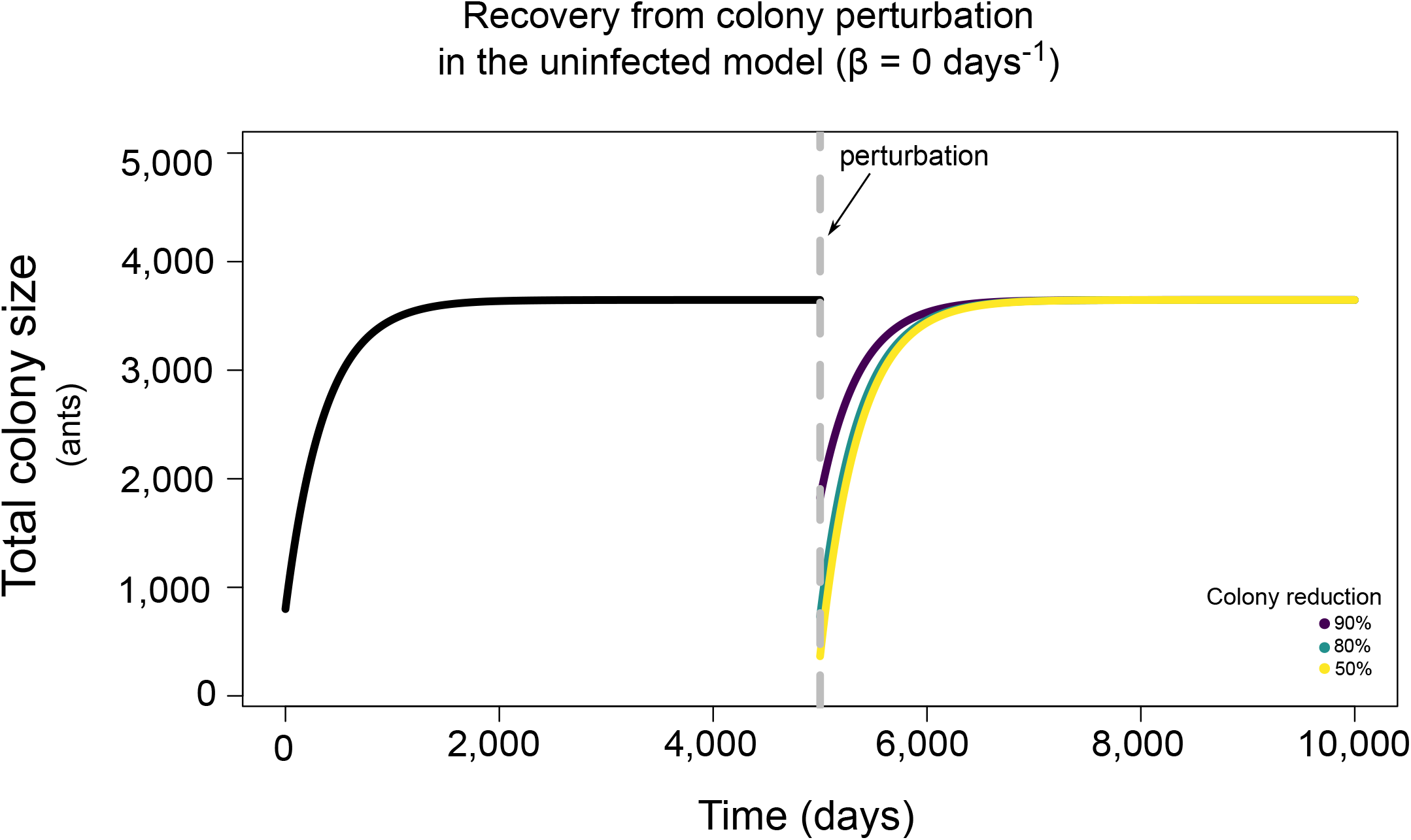
Recovery dynamics from the uninfected model in the face of perturbation. Recovery dynamics (from numerical simulation) for the uninfected model (eqs. 1-3) under baseline parameter values given in Table 1. Initial colony conditions were set at Fs = 200 ants, N = 500 ants, and B = 100 ants and the colony was allowed to go to equilibrium. At t = 5,000 days, the colony size was reduced by 50%, 80%, or 90% (applied across all compartments) and allowed to return to equilibrium.

**Fig. S2.**
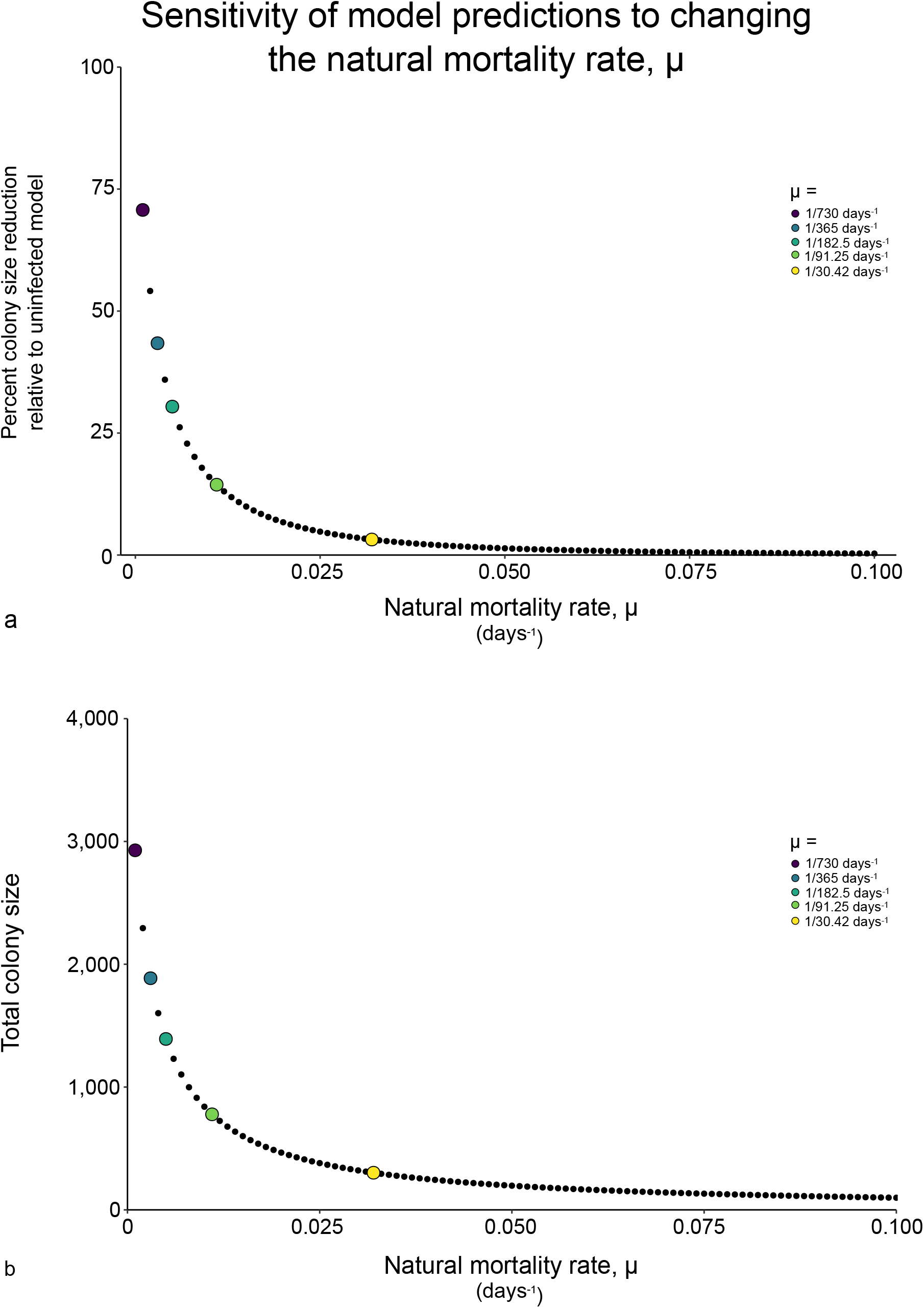
Sensitivity of model predictions to changing the natural mortality rate, *µ*. Model predictions are from numerical simulation under baseline parameter values given in Table 1. (a) The proportion reduction in colony size relative to the uninfected model under changing values of *µ* from 0 to 0.1 days^−1^. (b) Total colony size (Fs + N + B) under changing values of *µ* from 0 to 0.1 days^−1^.

**Fig. S3.**
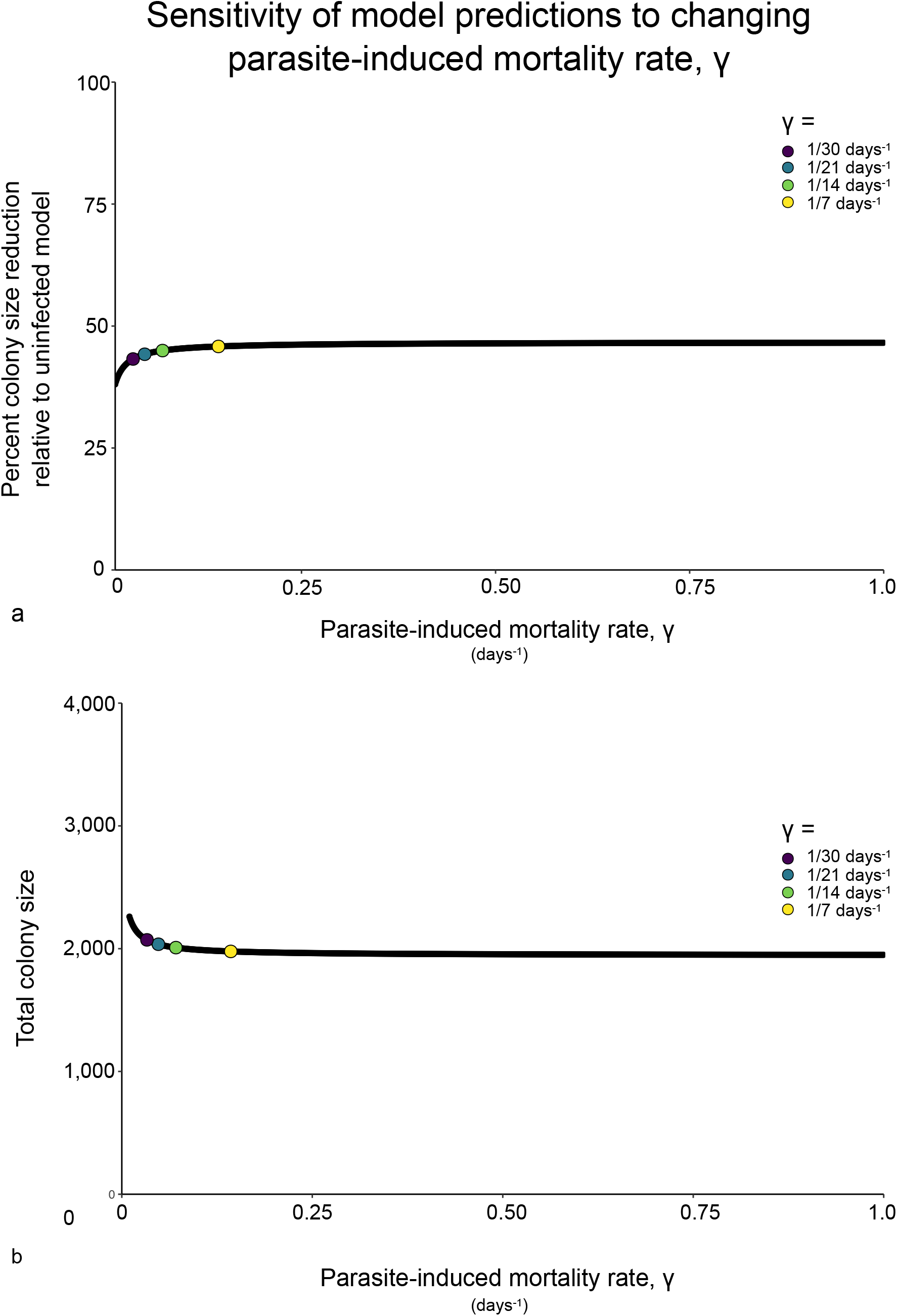
Sensitivity of model predictions to changing the parasite-induced mortality rate, *γ*. Model predictions are from numerical simulation under baseline parameter values given in Table 1. (a) The proportion reduction in colony size relative to the uninfected model under changing values of *γ* from 0 to 1 days^−1^. (b) Total colony size (Fs + N + B) under changing values of *γ* from 0 to 1 days^−1^.

**Fig. S4.**
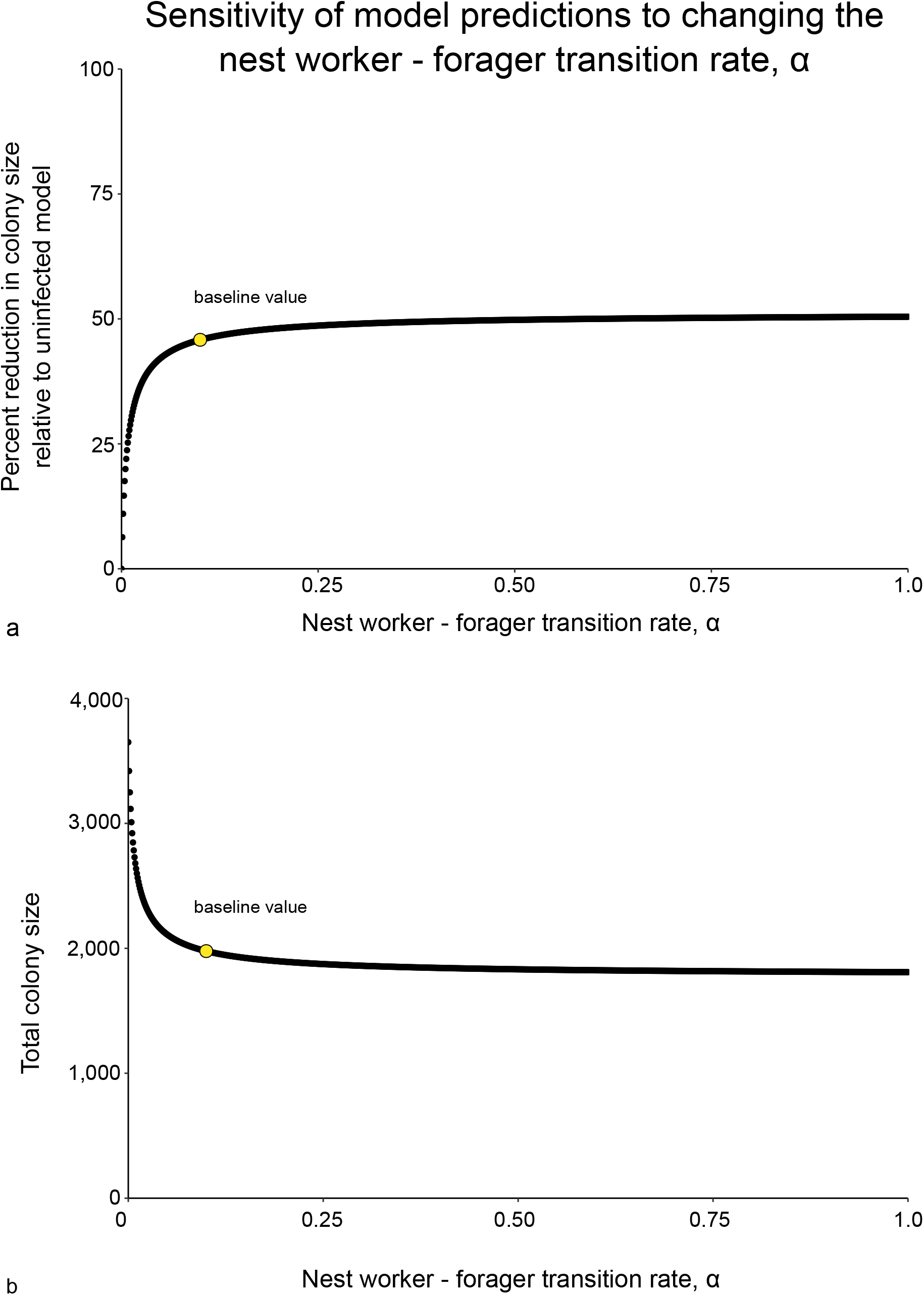
Sensitivity of model predictions to changing the nest worker – forager transition term, *α*. Model predictions are from numerical simulation under baseline parameter values given in Table 1. (a) The proportion reduction in colony size relative to the uninfected model under changing values of *α* from 0 to 1. (b) Total colony size (Fs + N + B) under changing values of *α* from 0 to 1.

**Fig. S5.**
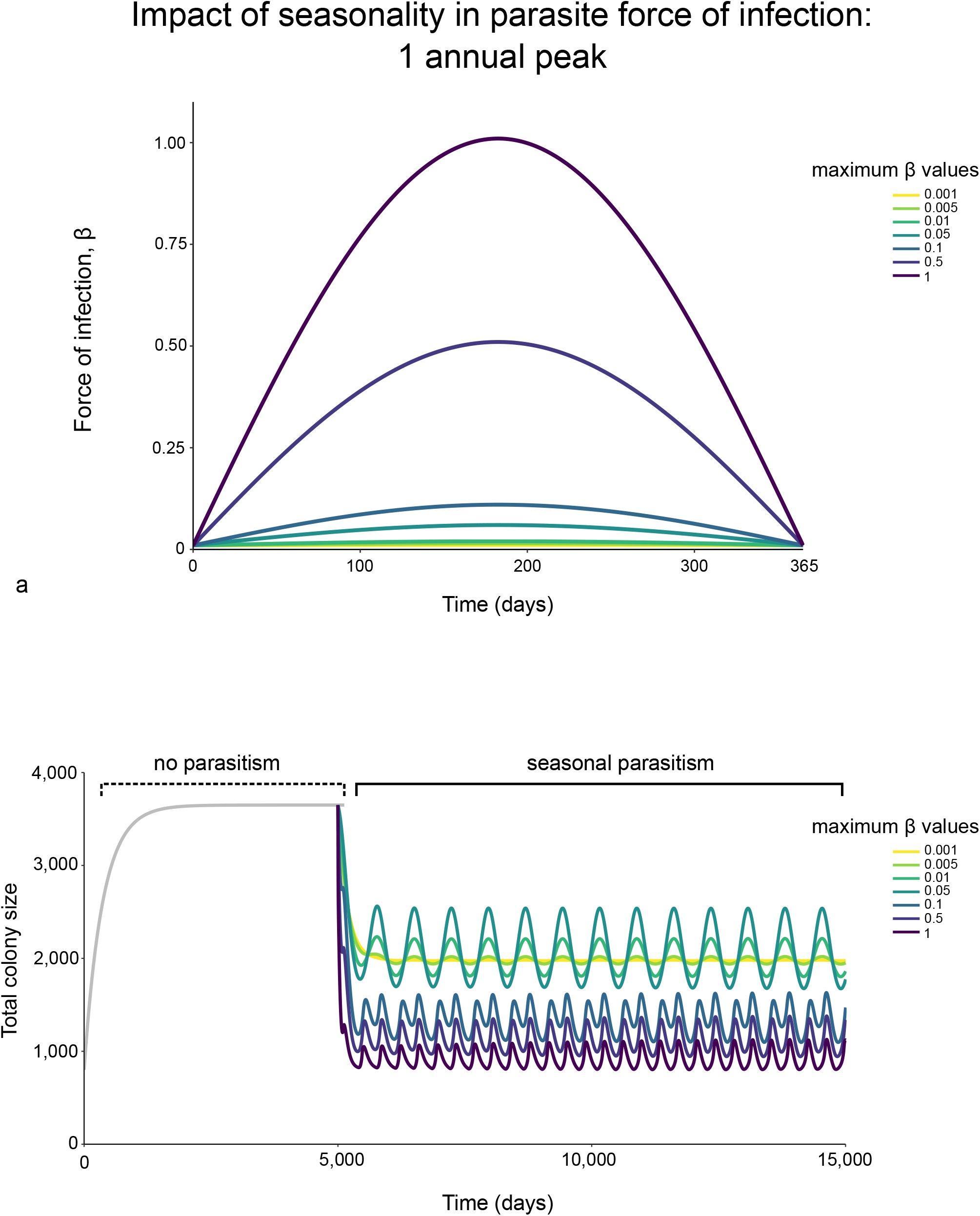
Impact of seasonality in parasite force of infection on model predictions (one annual peak). Model predictions are from numerical simulation under baseline parameter values given in Table 1. (a) The proportion reduction in colony size relative to the uninfected model under different maximum values of *β* with one annual peak. (b) Total colony size (Fs + N + B) under different maximum values of *β* with one annual peak.

**Fig. S6.**
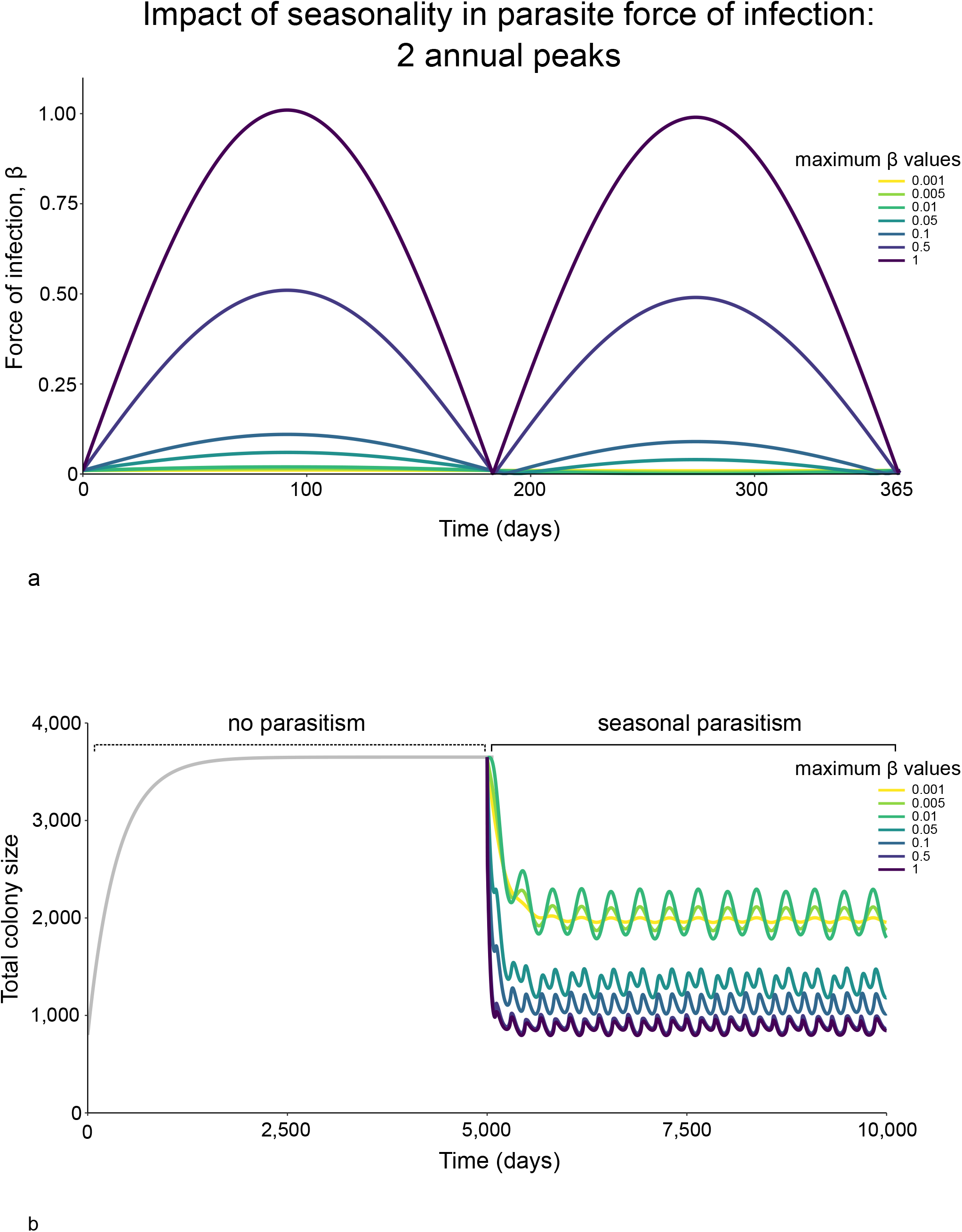
Impact of seasonality in parasite force of infection on model predictions (two annual peaks). Model predictions are from numerical simulation under baseline parameter values given in Table 1. (a) The proportion reduction in colony size relative to the uninfected model under different maximum values of *β* with two annual peaks. (b) Total colony size (Fs + N + B) under different maximum values of *β* with two annual peaks.

**Table S1.** Parameter estimates from the literature: colony birth rate

| Colony birth rate, $\lambda$ | | | |
| --- | --- | --- | --- |
| Ant species | Birth rate | Notes | Reference |
| *Solenopsis invicta* | 0 - 50 workers/day | Varies seasonally. | (97) |
| *Lasius neglectus* | 6 - 14 eggs/day/queen | - | (98) |
| *Solenopsis invicta* | 62 - 1776 eggs/day | Depends on the number of larvae already in the nest | (47) |
| *Leptothorax acervorum* | 1 - 23 eggs/day | - | (99) |
| *Dolichoderus mariae* | 27 - 40 eggs/day/queen | - | (100) |
| *Dolichoderus mariae* | 1500 - 6000 eggs/day/nest | - | (100) |
| *Iridomyrmex purpureus* | 93 - 175 eggs/day/queen | Depends on queen's rank dominance | (101) |

**Table S2.** Parameter estimates from the literature: brood developmental rate

| Brood developmental rate $1/\phi$ | | | | |
| --- | --- | --- | --- | --- |
| Ant species | Avg. time to adult emergence (days) | Range (days) | Reference | Notes |
| *Solenopsis invicta* | 18 | - | (102) | - |
| *Linepithema humile* | - | 40 - 140 | (103) | - |
| *Solenopsis invicta* | - | 23 - 55 | (104) | - |
| *Solenopsis invicta* | - | 20 - 31 | (105) | - |
| *Dinoponera quadriceps* | 95 | 90 - 100 | (106) | within (107) |
| *Myrmecia regularis* | - | 150 - 180 | (108) | within (107) |
| *Tetraponera anthracina* | 100 | - | (109) | within (107) |
| *Aenictus laeviceps* | 65 | - | (110) | within (107) |
| *Eciton burchelli* | 45 | - | (64) | within (107) |
| *Eciton hamatum* | 50 | - | (110) | within (107) |
| *Atta sexdens* | - | 40 - 60 | (111) | within (107) |
| *Messor aciculatus* | 69.3 | - | (112) | within (107) |
| *Messor pergandei* | 60 | - | (113) | within (107) |
| *Myrmica rubra* | 115 | - | (114) | within (107) |
| *Myrmica rubra* | 84 | - | (114) | within (107) |
| *Myrmica rubra* | 54 | - | (114) | within (107) |
| *Monomorium pharaonis* | 36.4 | - | (115) | within (107) |
| *Monomorium pharaonis* | - | 25 - 54 | (116) | within (107) |
| *Pogonomyrmex sp.* | - | 35 - 42 | (117) | within (107) |
| *Solenopsis invicta* | - | 24 - 25 | (118) | within (107) |
| *Solenopsis invicta* | - | 55 | (104) | within (107) |
| *Tetramorium caespitum* | - | 43 - 63 | (119) | within (107) |
| *Linepithema humile* | 44 | - | (120) | within (107) |
| *Wasmannia auropunctata* | - | 35 - 40 | (121) | within (107) |
| *Liometopum apiculatum* | 28.2 | - | (122) | within (107) |
| *Liometopum apiculatum* | 70.8 | - | (122) | within (107) |
| *Camponotus clariothorax* | 67 | - | (123) | within (107) |
| *Camponotus festinatus* | 69 | - | (123) | within (107) |
| *Camponotus laevigatus* | - | 48 - 70 | (123) | within (107) |
| *Camponotus modoc* | 55 | - | (123) | within (107) |
| *Camponotus planatus* | 57 | 54 - 58 | (123) | within (107) |
| *Camponotus sericeus* | - | 24 - 26 | (124) | within (107) |
| *Camponotus sericeus* | 55 | - | (124) | within (107) |
| *Camponotus sericeus* | 20 | - | (124) | within (107) |
| *Camponotus vicinus* | - | 54 - 70 | (123) | within (107) |
| *Formica polyctena* | - | 35 - 45 | (125) | within (107) |
| *Formica rufa* | - | 35 - 37 | (126) | within (107) |
| *Oecophylla longinoda* | 39 | - | (127) | within (107) |
| *Prenolepis imparis* | - | 70 - 90 | (128) | within (107) |

**Table S3.** Parameter estimates from the literature: proportion of the colony foraging

| Proportion of the colony that forages, $\kappa$ | | | |
| --- | --- | --- | --- |
| Ant species | % that are foragers | Notes | Reference |
| *Formica polystena* | 4.5 - 57.5% | - | (129) |
| *Solenopsis invicta* | 16 - 58% | - | (39) |
| *Pogonomyrmex owyheeii* | 16% | - | (130) |
| *Pogonomyrmex badius* | 5 - 42% | - | (42) |
| *Solenopsis invicta* | 10 - 90% | - | (131) |
| *Solenopsis invicta* | 30 - 80% | - | (131) |
| *Pogonomyrmex mendozanus* | 10 - 13% | - | (132) |
| *Pogonomyrmex inermis* | 15 - 44% | - | (132) |
| *Pogonomyrmex rastratus* | 7 - 10% | - | (132) |

**Table S4.** Parameter estimates from the literature: natural mortality rate

| Natural mortality rates, $\mu$, of workers (not reproductives) | | | |
| --- | --- | --- | --- |
| Ant species | Avg. life span (days) | Notes | Reference |
| *Atta colombica* | 3 - 4 days | Avg. lifespan for 50% surviving | (133) |
| *Cataglyphis bicolor* | 6 days | - | (134) |
| *Pogonomyrmex owyheeii* | 14 days | - | (130) |
| *Pogonomyrmex sp.* | 30 days | - | (135) |
| *Solenopsis invicta* | 60 - 540 days | - | (136) |
| *Solenopsis invicta* | 70 - 490 | - | (137) |
| *Oecophylla smaragdina* | 175 - 210 days | Avg. lifespan for 50% surviving | (138) |
| *Aphaenogaster rudis* | > 1095 days | Maximum recorded. | (139), within (10). |
| *Leptothorax lichtensteini* | 912 days | - | (140), within (10). |
| *Leptothorax nylanderi* | 1095 days | - | (140), within (10). |
| *Monomorium pharaonis* | 63 - 70 days | - | (115), within (10). |
| *Myrmecia gulosa* | 620 days | - | (141), within (10). |
| *Myrmecia nigriceps* | 803 days | - | (141), within (10). |
| *Myrmecia nigrocinta* | 438 days | - | (141), within (10). |
| *Myrmecia pilosula* | 474 days | - | (141), within (10). |
| *Myrmecia vindex* | 693 days | - | (141), within (10). |
| *Rhytidoponera purpurea* | 949 days | - | (141), within (10). |
| *Platythyrea punctata* | 100 - 200 days | - | (142) |
| *Lasius niger* | 325 days | Avg. lifespan for 50% surviving, control treatment | (143) |
| *Diacamma rugosum* | 208.8 days | - | (144) |

**Table S5.** Parameter estimates from the literature: parasite-induced mortality rates

| Parasite species | Parasite-induced mortality rate, $\gamma$ | | | Reference |
| --- | --- | --- | --- | --- |
|  | Trans. loca-<br>tion | Avg. development<br>time | Notes |  |
| *Ophiocordyceps unilateralis* | Ex-nido | 6 - 20 days | - | (31) |
| *Beauveria bassiana* | In-nido | 3 - 7 days | - | (71) |
| *Metarhizium anisopliae* | In-nido | 10 - 30 days | Varies based on colony genetic diversity: polygynous vs. monogynous | (145) |
| *Metarhizium anisopliae* | In-nido | 3 - 9 days | - | (89) |
| *Pseudacteon litoralis* | Ex-nido | 46 days | For larval + pupal development inside ant | (44) |
| *Pseudacteon litoralis* | Ex-nido | 30 - 49 days | For larval + pupal development inside ant | (146) |
| *Pseudacteon tricuspis* | Ex-nido | 23 - 50 days | For larval + pupal development inside ant | (146) |
| *Pseudacteon browni* | Ex-nido | 23 - 43 days | For larval + pupal development inside ant | (146) |
| *Pseudacteon crawfordi* | Ex-nido | 24 - 32 days | For larval + pupal development inside ant | (146) |

**Table S6.** Parameter estimates from the literature: foraging rates

| Ant species | Foraging rate, a component of $\beta$ | | Reference |
| --- | --- | --- | --- |
|  | Avg. foraging<br>trips/day per ant | Notes |  |
| *Pogonomyrmex barbatus* | 1 - 25 (24 hrs) | - | (82) |
| *Paraponera clavata* | 1 - 9 (12 hrs) | - | (80) |
| *Cataglyphis bicolor* | 3.7 (24 hrs) | - | (81) |
| *Pogonomyrmex montanus* | 50 - 950 trips/day per colony | Reported as mean foraging trips per day for entire colonies, not for individual foragers. Number of foraging trips/day/colony varies seasonally. | (117) |
| *Pogonomyrmex subnitidus* | 50 - 10,500 trips/day per colony | Reported as mean foraging trips per day for entire colonies, not for individual foragers. Number of foraging trips/day/colony varies seasonally. | (117) |
| *Pogonomyrmex rugosus* | 50 - 25,000 trips/day per colony | Reported as mean foraging trips per day for entire colonies, not for individual foragers. Number of foraging trips/day/colony varies seasonally. | (117) |

**Table S7.** Parameter estimates from the literature: parasite encounter rates

| Parasite encounter rate, a component of $\beta$ | | | |
| --- | --- | --- | --- |
| Ant species | Parasite | Attack rate | Reference |
| *Linepithema sp.* | *Pseudacteon sp.* | No direct attack rates provided, but see results section. | (84) |
| *Azteca instabilis* | *Pseudacteon sp.* | 0 - 8 attacks/15 mins | (147) |
| *Atta vollenweideri*, *Acromyrmex sp.* | Various | 0.3 - 1.2 attacks/minute | (83) |

